# The representational dynamics of visual objects in rapid serial visual processing streams

**DOI:** 10.1101/394148

**Authors:** Tijl Grootswagers, Amanda K. Robinson, Thomas A. Carlson

## Abstract

In our daily lives, we are bombarded with a stream of rapidly changing visual input. Humans have the remarkable capacity to detect and identify objects in fast-changing scenes. Yet, when studying brain representations, stimuli are generally presented in isolation. Here, we studied the dynamics of human vision using a combination of fast stimulus presentation rates, electroencephalography and multivariate decoding analyses. Using a presentation rate of 5 images per second, we obtained the representational structure of a large number of stimuli, and showed the emerging abstract categorical organisation of this structure. Furthermore, we could separate the temporal dynamics of perceptual processing from higher-level target selection effects. In a second experiment, we used the same paradigm at 20Hz to show that shorter image presentation limits the categorical abstraction of object representations. Our results show that applying multivariate pattern analysis to every image in rapid serial visual processing streams has unprecedented potential for studying the temporal dynamics of the structure of representations in the human visual system.

## Introduction

The human brain can effortlessly extract abstract meaning, such as categorical object information, from a visual image, and can do so in less than 200 milliseconds (Carlson, Tovar, Alink, & Kriegeskorte, 2013; Cichy, Pantazis, & Oliva, 2014; Contini, Wardle, & Carlson, 2017; Keysers, Xiao, Földiák, & Perrett, 2001; Mack, Gauthier, Sadr, & Palmeri, 2008; Mack & Palmeri, 2011; Potter, 1975, 1976; Potter, Wyble, Hagmann, & McCourt, 2014; VanRullen & Thorpe, 2001). The temporal dynamics of the emerging representation of visual objects has been studied extensively using multivariate decoding methods and neuroimaging methods with high temporal resolution, such as EEG and MEG. In these experiments, stimuli are generally presented with a large inter-stimulus interval (ISI) to avoid contamination from temporally adjacent stimuli, typically around one second (Carlson et al., 2013; Cichy et al., 2014; Grootswagers, Ritchie, Wardle, Heathcote, & Carlson, 2017; Isik, Meyers, Leibo, & Poggio, 2014; Kaneshiro, Guimaraes, Kim, Norcia, & Suppes, 2015). This design allows the brain to process each stimulus and avoids temporally overlapping stimulus representations. While such designs have yielded important insights into the representational dynamics of object processing, in the natural world, we are bombarded with a constant stream of changing visual input. The standard paradigm, in which stimuli are presented in isolation with a large ISI, thus may not yield the most accurate description the temporal dynamics of emerging object representations in the real world. One major advantage of multivariate decoding methods (Grootswagers, Wardle, & Carlson, 2017; Haynes, 2015) is that they allow testing for statistical dependencies in data without a resting baseline. Exploring representational dynamics using decoding and fast visual presentation rates therefore offers unique potential for investigating visual processing.

Here, we diverge from the traditional approach and propose a new method for studying the representational dynamics of human vision. It has been shown previously that stimuli presented at high presentation rates are all processed to some degree by the visual system and that their neural representations can co-exist in the visual system (Marti & Dehaene, 2017; Mohsenzadeh, Qin, Cichy, & Pantazis, 2018; Rossion, Torfs, Jacques, & Liu-Shuang, 2015; Rousselet, Fabre-Thorpe, & Thorpe, 2002). Behavioural work has additionally shown that the human visual system can extract abstract information from a visual stimulus at very fast presentation rates (Crouzet, Kirchner, & Thorpe, 2010; Keysers et al., 2001; Macé, Thorpe, & Fabre-Thorpe, 2005; Mack et al., 2008; Mack & Palmeri, 2015; Marti & Dehaene, 2017; Potter, 1975, 1976; Potter et al., 2014; Rossion et al., 2015; Thorpe, Fize, & Marlot, 1996). In the current study, we draw on this human capacity and study visual object recognition using fast stimulus presentation rates and multivariate decoding analyses of EEG evoked responses (Grootswagers, Wardle, et al., 2017). We used a rapid serial visual presentation (RSVP) paradigm to study the representations of a large set of 200 visual objects presented at a speed of 5 images per second (5Hz; 200ms per image). The objects were carefully selected to allow categorisation at three different levels of abstraction. The high presentation rate enabled us to obtain 40 repetitions of 200 different stimuli in a short EEG session. The increased power elicited by the faster image presentation rates allowed us to use a much larger stimulus set than previous studies, and to analyse neural responses to all distractors, rather than a single target, in the stream. We additionally examined the effect of higher level cognitive processes on the emerging representations by having participants detect targets that were identifiable based on low-level visual features or abstract categories in separate trials. In doing so, we could disentangle the temporal dynamics of visual processing and categorical abstraction of non-target stimuli from target selection processes. We successfully decoded different categorical contrasts for the 200 objects, suggesting that individual stimuli were processed up to abstract categorical representations. Strikingly, we found similar results in a follow-up experimental session, where we used a much higher presentation rate of 20 images per second (20Hz; 50ms per image). The unprecedented ability to test such large numbers of different stimuli in relatively short EEG scanning sessions shows great potential for studying the dynamics of the structure of information in the human visual system.

## Methods

All stimuli and data can be found at https://doi.org/10.17605/OSF.IO/A7KNV.

### Stimuli

We collected a stimulus set of 200 visual objects from different categories. Stimuli were obtained from the free image hosting website www.pngimg.com. The categories were manually selected, guided by categorical hierarchies described in the literature (Caramazza & Mahon, 2003; Caramazza & Shelton, 1998; Carlson et al., 2013; Connolly et al., 2012; Grill-Spector & Weiner, 2014; Kiani, Esteky, Mirpour, & Tanaka, 2007; Kriegeskorte, Mur, Ruff, et al., 2008; Mahon & Caramazza, 2011; Peelen & Caramazza, 2012; E. H. Rosch, 1973). There were two high level categories (animate, inanimate) consisting of 10 categories (5 animate, and 5 inanimate categories). Each of these 10 categories (e.g., mammal, tool, flower) was further separated into 5 object categories (e.g., cow, dog, giraffe, etc.), which consisted of 4 images each (Figure 1a). During the experiment, participants were instructed to count target stimuli (Figure 1b). To examine how attending to different features of the stimuli affected the emerging representations, we used two different sets of target stimuli. The target stimuli were either boats, or geometric star shapes, and there were eight exemplars of each target type (Figure 1 – inset). We hypothesized that detecting the star shapes among the other objects was possible using low level visual cues, while for recognising boat targets, it was necessary to process stimuli to a more abstract categorical level.

**Figure 1.**
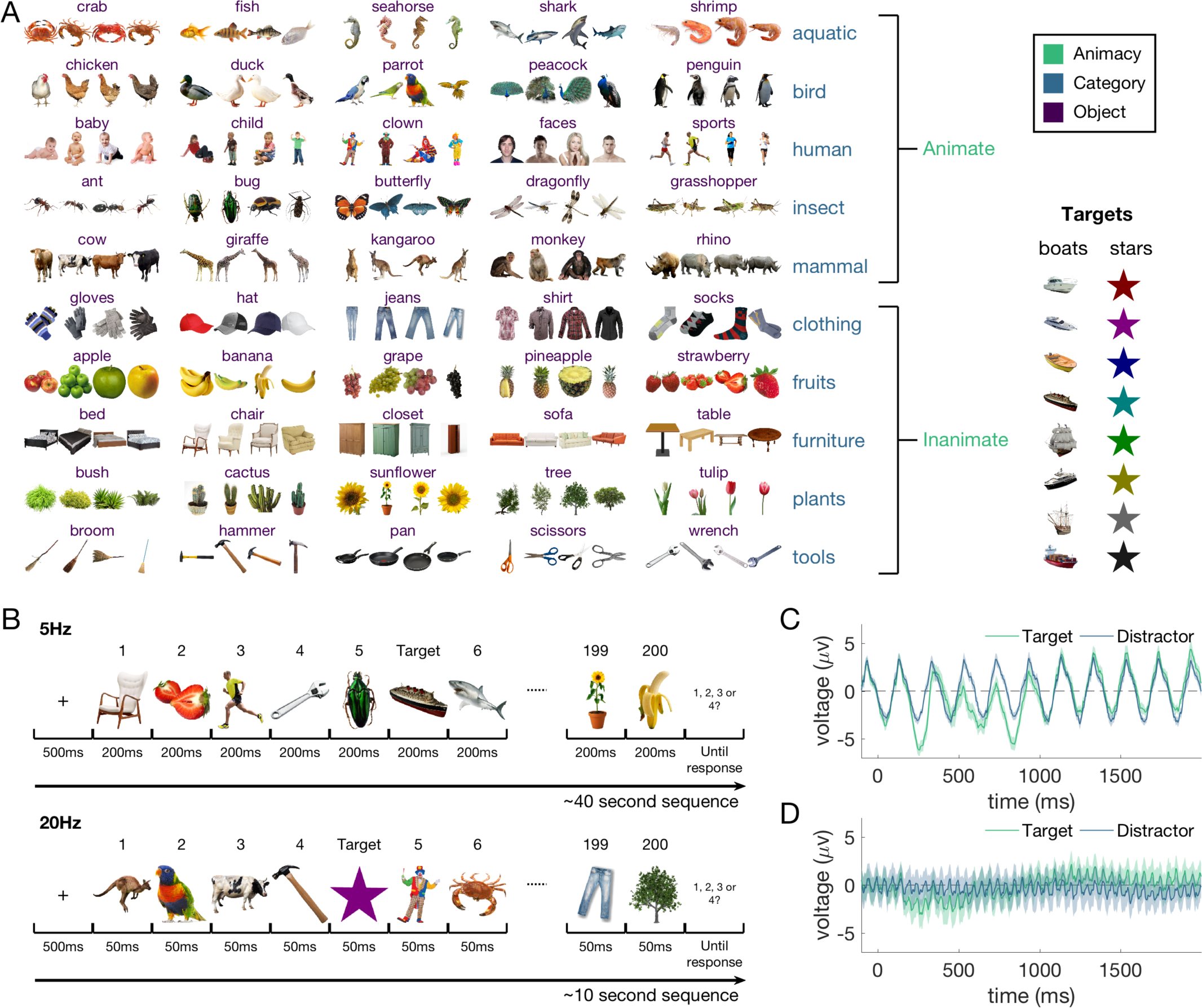
Stimuli and design. A) Experimental stimuli. There were 200 images of objects (obtained from www.pngimg.com), organised in categories at three different levels: Animacy (animate, inanimate), category (10 categories e.g., mammal, tool, flower) and object (50 categories e.g., cow, dog, giraffe). In the experiment, participants were asked to count the number of target objects from two categories: boats and geometric star shapes, each with eight images. B) Experimental design. Trials consisted of all 200 images presented in random order, with 1-4 targets interspersed throughout. Images were presented in 5Hz sequences (200ms each) in session 1, and 20Hz sequences (50ms each) in session 2. C,D) Subject-averaged event-related potentials (ERPs) at channel Oz for target and non-target images in the 5Hz (C) and 20Hz (D) sequences (shaded areas show the standard-error across subjects).

### Participants and experimental procedure

Participants were 16 adults recruited from the University of Sydney (5 females; age range 18-38 years) in return for payment or course credit. The study was approved by the University of Sydney ethics committee and informed consent was obtained from all participants. Participants viewed 40 sequences of objects, each lasting between 40.2 -40.8 seconds (depending on the number of targets in the sequence). In each sequence, the 200 stimuli were presented in random order, for a duration of 200ms each with no gap between successive images (5Hz). In addition to the 200 stimuli, target stimuli were inserted throughout the sequence (Figure 1b). In half of the sequences, the target stimuli were boats, and in the other sequences, the target stimuli were geometric stars (Figure 1). A random number between 1 and 4 targets were presented in the sequence, with the condition that targets could not appear within the first 10 or last 10 images, and ensuring there were at least 12 non-target stimuli between subsequent targets. At the start of each sequence, participants were prompted to count the number of targets in the sequence (“Count the boats in the trial” or “Count the stars in the trial” in random order) and the 8 potential targets were shown. They were instructed to respond at the end of the sequence using a 4-way button box. After each sequence, participants received feedback. They started the next sequence with a button press. This session lasted approximately 40 minutes in total. After a short break, the second experimental session started, and participants performed another 40 sequences using the same procedure as session one, except that the images were presented for only 50ms (a presentation speed of 20Hz). The second session lasted about 10 minutes.

### EEG recordings and preprocessing

Continuous EEG data were recorded using a BrainVision ActiChamp system, digitized at a 1000-Hz sample rate. The 64 electrodes were arranged according to the international standard 10–10 system for electrode placement (Oostenveld & Praamstra, 2001). During recording, all scalp electrodes were referenced to Cz. Preprocessing was performed offline using EEGlab (Delorme & Makeig, 2004). Data were filtered using a Hamming windowed FIR filter with 0.1 Hz highpass and 100Hz lowpass filters, and were downsampled to 250Hz. No further preprocessing steps were applied, and the channel voltages at each time point were used for the remainder of the analysis. Epochs were created for each stimulus presentation (except targets) ranging from [-100 to 1000ms] relative to stimulus onset. We initially had used the same range for target-distractor decoding but found that this window did not capture the full process. Therefore, for comparing targets versus distractors, we created larger epochs ranging from [-100 to 2000ms] relative to the onset of a target. For each target t, we selected at random another distractor in the same sequence and created a matching epoch relative to the onset of that distractor. Choosing distractors in this way meant that the number of targets and distractors were balanced and matched per sequence (and chance level accuracy is 50%) and that the neural representations of targets and distractors were unlikely to overlap in a consistent manner. Event-related potentials (Figure 1C&D) for both the targets and non-targets exhibited clear signal at the presentation frequencies (see Figure S1 for the associated scalp maps and amplitude spectra).

### Decoding analysis

We applied an MVPA decoding pipeline (Grootswagers, Wardle, et al., 2017; Oosterhof, Connolly, & Haxby, 2016) to the EEG channel voltages, consisting of a regularised linear discriminant analysis (LDA) classifier applied in an exemplar-by-sequence-cross-validation approach. Decoding was performed within subject, and the results were analysed at the group level. This pipeline was applied to each stimulus presentation epoch in the sequence to investigate object representations in fast sequences. To investigate the temporal dynamics of target selection, we compared neural responses to targets with those to non-target distractor stimuli. Classifiers were then trained to distinguish targets from non-targets separately for the 5Hz and 20Hz sequences, and for boat and star target sequences.

We investigated object representations for the 200 non-target images using multiple categorical distinctions. First, we decoded three contrasts that impose different amounts of categorical abstraction. At the highest level, we decoded animacy (i.e., animate versus inanimate objects). The next contrast was the category tier (10 classes, e.g., mammal, insect, furniture, tool, etc.) where we decoded all 45 possible pairwise combinations. The lowest categorical level was the object level (50 classes, e.g., cow, butterfly, table, hammer, etc.). Here, we decoded all 1225 possible pairwise object combinations (i.e., cow versus butterfly, cow versus table, etc.). Finally, at the lowest level, we investigated image-level representations by decoding all 19900 possible pairwise combinations of the 200 stimuli. We report the mean pairwise classification accuracies, so that chance-level accuracy for all comparisons is at 50%, which aids comparing accuracies across contrasts.

To investigate similarities in underlying object representation signals between the 5Hz and 20Hz presentations, we used a temporal generalisation approach (Carlson, Hogendoorn, Kanai, Mesik, & Turret, 2011; King & Dehaene, 2014; Meyers, Freedman, Kreiman, Miller, & Poggio, 2008). To test generalisation between the conditions, we trained classifiers on all time points in the data from the 5Hz sequences and tested them on all time points in the data from the 20Hz sequences. We repeated this for the inverse (training on 20Hz and testing on 5Hz), and averaged the resulting time-generalisation matrices (Kaiser, Azzalini, & Peelen, 2016).

All steps in the decoding analysis were implemented in CoSMoMVPA (Oosterhof et al., 2016). For the categorical contrasts that grouped more than one image, we used an image-by-sequence-cross-validation scheme so that identical images were not part of both training and test set (Carlson et al., 2013; Grootswagers, Wardle, et al., 2017). This was implemented by first splitting the data into four sets, where the first set consisted of the first images from each of the 50 object categories (i.e., cow-1, table-1 etc.), the second set of the second images (i.e., cow-2, table-2 etc.), etc. One of these sets was used as test data, and the other three as training data for the leave-one-sequence out cross-validation, where all data from one sequence was used as test data, and data from the remaining sequences as training data. For each decoding contrast, this resulted in 160 (4 images by 40 sequences) cross-validation partitions. Where image-by-sequence cross-validation was not possible (i.e., image-level and target-distractor decoding), we used a leave-one-sequence-out cross-validation scheme, where all epochs from one sequence were used as test set, resulting in 40 cross-validation partitions. We used a linear discriminant analysis (LDA) classifier (implemented in CoSMoMVPA) and report the mean cross-validated decoding accuracy.

### Representational Similarity Analysis

To study the emerging representational structure of our 200 stimuli, we analysed our data using the Representational Similarity Analysis (RSA) framework (Kriegeskorte & Kievit, 2013; Kriegeskorte, Mur, & Bandettini, 2008; Kriegeskorte, Mur, Ruff, et al., 2008), which allows comparing models of object representations. The decoding results at the image level were organised into a 200 by 200 neural representational dissimilarity matrix (RDM), which for each pair of images, contained the mean cross-validated decoding accuracy (images that evoke more dissimilar neural responses are better decodable). One neural RDM was created for each subject, and each time point (group mean RDM at 100-400ms shown in Figure 2, top row). We compared the neural RDMs to six candidate models; first, we created one model for each of the three categorical levels, grouping images from the same category (Figure 2, second row). We also used three low-level image feature control models (Figure 2, third row), which were created by correlating the vectorized experimental images. The models consisted of an image silhouette similarity model, which is based on the binary alpha layer of the stimuli and is a good predictor of differences in brain responses (Carlson et al., 2011; Teichmann, Grootswagers, Carlson, & Rich, 2018; Wardle, Kriegeskorte, Grootswagers, Khaligh-Razavi, & Carlson, 2016)), a model based on the CIELAB-colour values of the stimuli, and a model based on the difference in luminance of the stimuli. Figure 2 shows the candidate models and the correlation distance between each of the candidate models (bottom row). The small correlations between the categorical models and the low-level feature models suggests that there was little overlap between the low-level features and categorical organisations in the stimulus set. To quantify the unique contributions of all models to the neural dissimilarities, we modelled the time-varying neural RDMs of each subject as a linear combination of the candidate models using a GLM (Oosterhof et al., 2016; Proklova, Kaiser, & Peelen, 2017); for each time point, the lower triangles of the neural RDM and candidate models were vectorised, and regression coefficients were obtained for all candidate models. This resulted in one beta estimate for each model, subject, and time point. We then analysed at the group level the mean beta estimates across subjects. To visualise the dynamic representational structure, at each point in time, we created a two-dimensional embedding of all 200 images. To compute the two-dimensional embedding, we applied t-SNE (Maaten & Hinton, 2008) to the mean neural RDMs. This approach finds an embedding of the multi-dimensional space in a two-dimensional representation so that the distances between points reflect their multidimensional pattern dissimilarities as best as possible.

**Figure 2.**
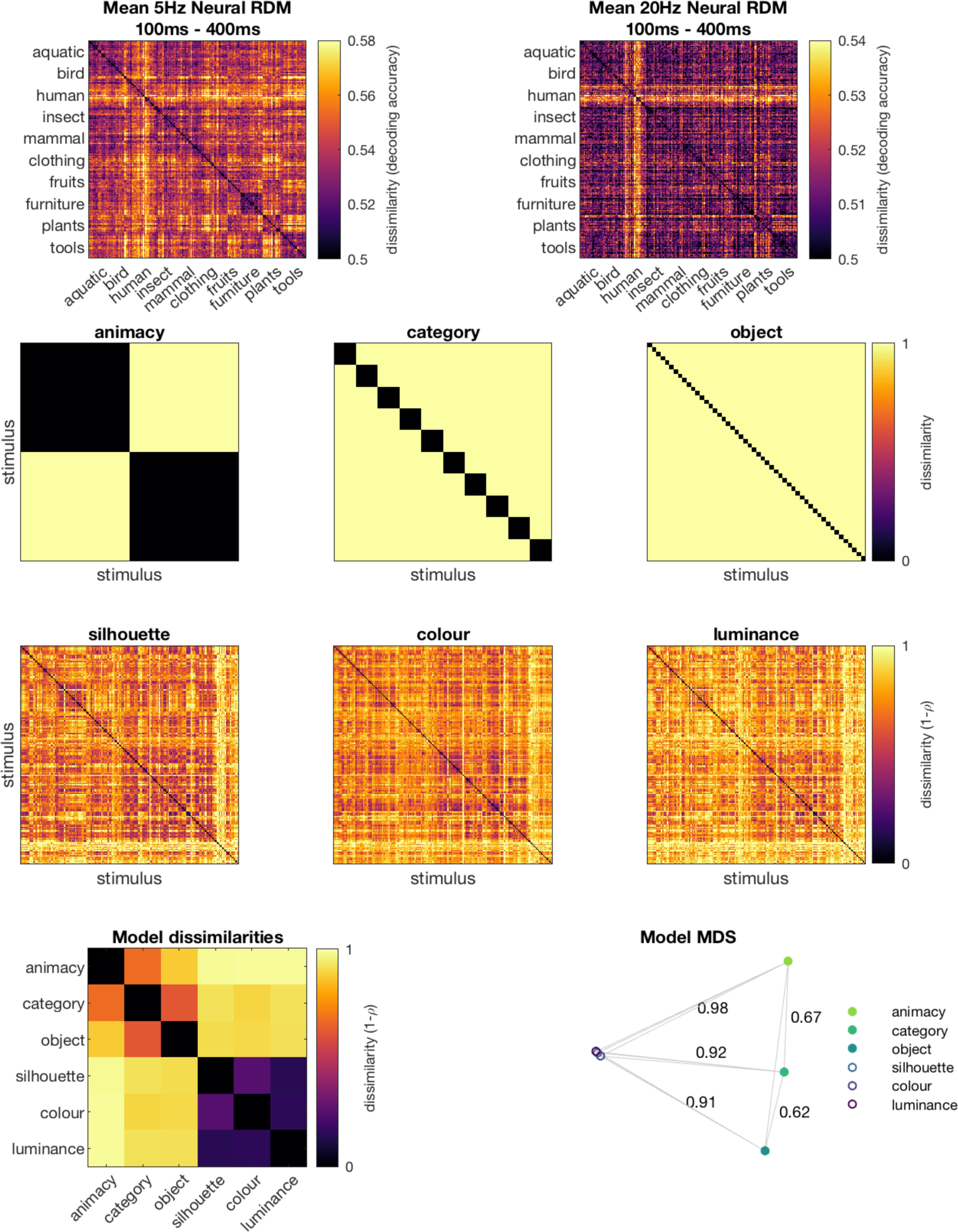
Candidate models used in the RSA. Top row: time-averaged neural RDMs for the 5Hz and 20Hz conditions. Each point in the 200 by 200 matrix represents the dissimilarity (here: decoding accuracy) between a pair of images. Second row: categorical models predict that responses to stimuli from the same category are more similar than responses to stimuli for different categories. Third row: image properties entered the regression as control models to quantify the contribution of low-level visual differences to the neural dissimilarities. Bottom row left: dissimilarities (1-correlation) between all candidate models. The order of the images in the 200 x 200 RDMs are the same as in Figure 1. Bottom row right: model dissimilarities projected in a 2-dimensional space using classical multi-dimensional scaling, which returns a configuration so that the distance between points approximates their dissimilarities. Annotated are the dissimilarities (1-correlation) between category-animacy and category-object, and between the silhouette model and all three categorical models.

### Statistical inference

In this study, we used Bayes factors (Dienes, 2011; Jeffreys, 1998; Rouder, Speckman, Sun, Morey, & Iverson, 2009; Wagenmakers, 2007) to determine the evidence for the null and alternative hypotheses. For the alternative hypothesis of above-chance decoding or correlation, a uniform prior was used ranging from the maximum value observed during the baseline (before stimulus onset) up to 1 (e.g., 100% decoding). For testing a non-zero difference between decoding accuracies, a uniform prior was used ranging from the maximum absolute difference observed during the baseline up to 50% (0.5). We then calculated the Bayes factor (BF) which is the probability of the data under the alternative hypothesis relative to the null hypothesis. We thresholded BF>3 and BF>10 as substantial and strong evidence for the alternative hypothesis, and BF<1/3 and BF<1/10 for substantial/strong evidence in favour of the null hypothesis (Jeffreys, 1998; Wetzels et al., 2011). BF that lie between 1/3 and 3 indicate insufficient evidence for either hypothesis.

## Results

We examined the representational dynamics of 200 different visual objects (Figure 1A), presented in 5Hz and 20Hz sequences (Figure 1B) using EEG. During the sequences, participants detected targets (boats or stars).

### The effect of target type and target selection

Participants were generally above chance (25%) at detecting targets (boats or stars) in the 5Hz and 20Hz sequences (Figure 3A-B). There was no difference in performance between the boat and star conditions (all BF < 1/3). On incorrect trials, responses often differed no more than one from the correct answer (Figure 3, right columns). This indicates that in general, participants missed at most one target when they responded incorrectly.

**Figure 3:**
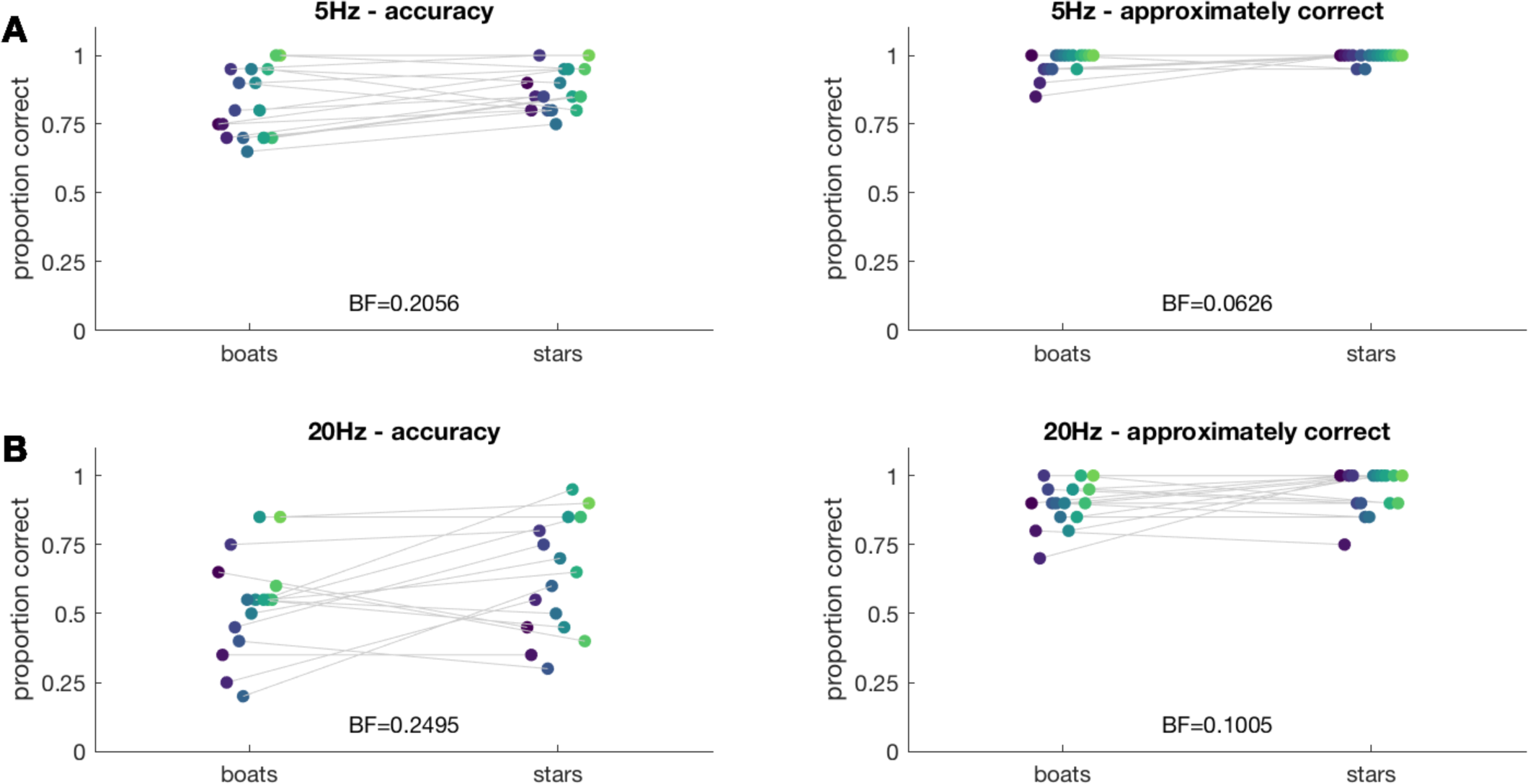
Behavioural results of target detection performance. (A) 5Hz sequences. (B) 20Hz sequences. Left columns show the mean proportion of correct responses for each participant separately for boat target sequences and star target sequences. Right columns show the mean approximately-correct (i.e., response differed by at most 1 from the correct answer) accuracy for each participant. Bayes Factors (BF) comparing mean accuracies between the boat and star sequences are listed above the x-axis.

The temporal dynamics of target selection were revealed by decoding targets from non-targets. The time-varying mean target-distractor decoding accuracy was computed separately for boat sequences and star sequences (Figure 4). Target-distractor decoding performance peaked around 67% in the 5Hz condition (Figure 4A), and around 60% in the 20Hz condition (Figure 4B). For both presentation rates, peak decoding performance was around 500ms. In both conditions, decoding for star targets was above chance earlier than for boats, which suggests that stars targets were easier to distinguish overall. Decoding performance remained above chance for over 1000 ms in the 5Hz sequences, and for approximately 800 ms in the 20Hz sequences.

**Figure 4.**
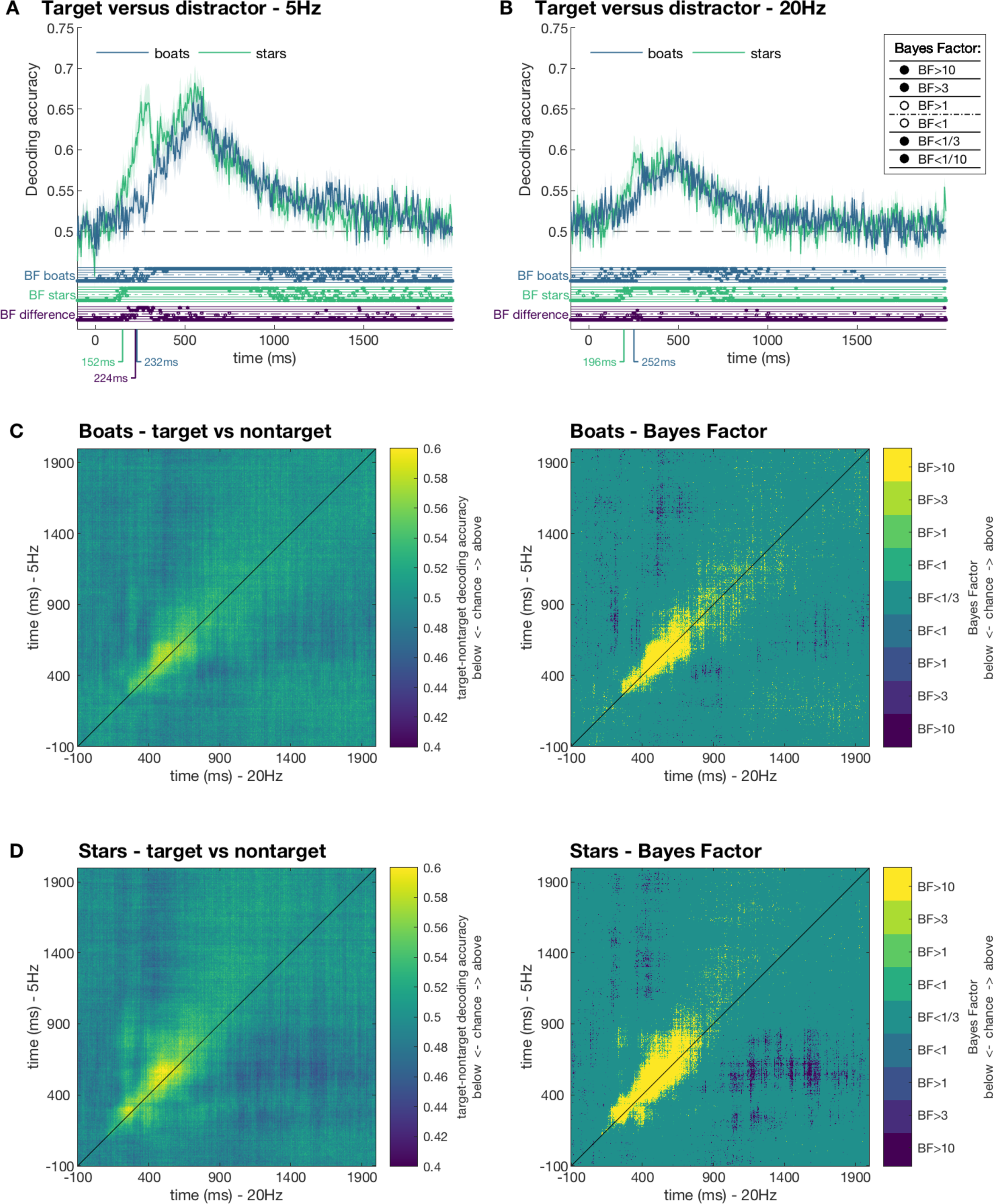
Decoding target versus distractor. For each target, a distractor was randomly selected from the same sequence, and classifiers were trained on target versus distractor. A-B) Plots show the mean leave-one-sequence-out cross-validated accuracy for the 5Hz condition (A), and the 20Hz condition (B). Shaded areas show the standard error of the mean across participants. Results are shown separately for boat target sequences and star target sequences. Dots below plots indicate thresholded Bayes Factors (BF, see inset) for the boat (top row) and star (middle row) sequences compared to chance and for the difference between boat and star sequences (bottom row). Annotated below the x-axis are the time points where the BF first exceeded 3 (for at least 2 consecutive time points). C-D) temporal generalisation results. The left columns show classifier generalisation performance for the boat (C) and star (D) between the different presentation durations. The right columns show corresponding thresholded Bayes Factors (yellow indicating above chance, and blue indicating below chance decoding). Higher than chance generalisation (yellow) on the diagonal indicates similar temporal dynamics of processing in the 5Hz condition as the 20Hz condition.

The temporal generalisation approach revealed target selection was very similar between the 5Hz and 20Hz sequences. For both boat and star target sequences, the onset of target decoding occurred around the same time, and cross-decoding was most evident along the diagonal, suggesting that target selection processes occurred at the same latencies regardless of the sequence speed and image duration (Figure 4c-d).

### Decoding categorical contrasts of 200 stimuli

In the 5Hz condition, we observed above chance decoding for all categorical levels (Figure 5, blue lines), starting at 100ms after stimulus onset for the categorical levels, and earlier (80ms) at the image level. This difference may be caused by decodable low-level visual features at the image level, which are controlled for by the exemplar-cross-validation approach at the categorical levels (Carlson et al., 2013; Grootswagers, Wardle, et al., 2017). These decoding onsets correspond well to the existing decoding literature, which has reported onsets for various categories between 80ms and 100ms (Carlson et al., 2013; Cichy et al., 2014; Kaneshiro et al., 2015). For the animacy level, the results showed three distinct peaks in decoding performance (150, 200ms and 400ms). In contrast, peak decoding happened around 200ms for category and object decoding and 130ms for image decoding. For all categorical levels, above-chance decoding was sustained until around 500ms. Note that at 500ms, there were already two new stimuli presented.

**Figure 5.**
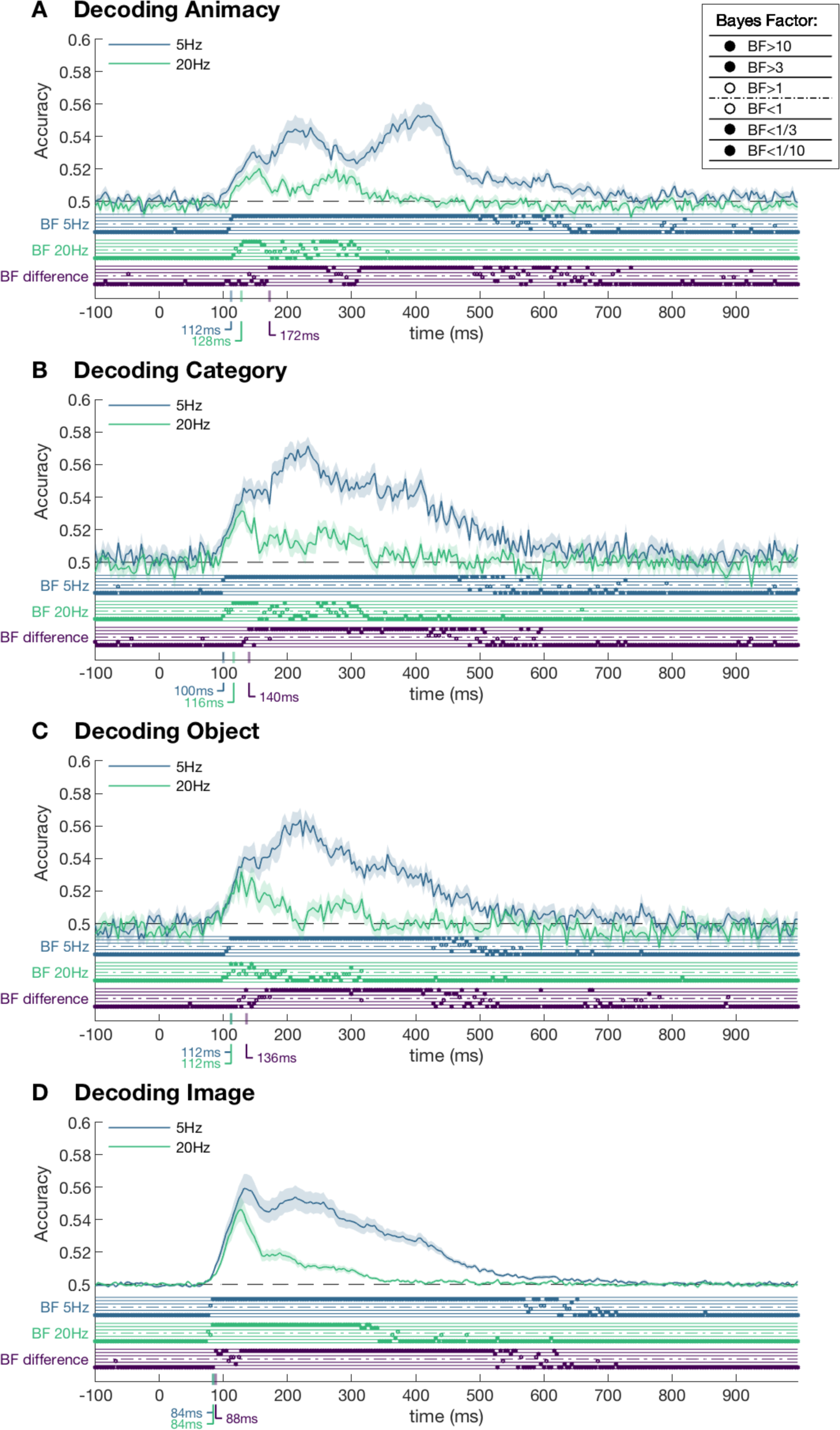
Mean decoding accuracy for 5Hz and 20Hz conditions. A) Decoding animacy (animate versus inanimate). B) Mean pairwise decoding for the 10 categories (e.g., mammal, tools). C) Mean pairwise decoding for 50 object categories (e.g., dog, giraffe). D) Mean pairwise decoding for all 200 images. Shaded areas depict standard error of the mean across subjects. Dots below plots indicate thresholded Bayes Factors (BF, see inset) for the 5Hz condition compared to chance (top rows), 20Hz condition compared to chance (middle rows) and for the difference between the 5Hz and 20Hz results (bottom rows). The time points where the BF first exceeded 3 are annotated below the x-axis.

In the 20Hz condition (Figure 5, green lines), we again observed above-chance decoding for all levels. Notably, the onset of decoding was around the same time point as in the 5Hz condition and subsequent decoding followed the same trajectory but diverged later in the time series (indicated by the bottom row of Bayes factors). The overall peak decoding performance was lower, and the peak decoding time points appeared earlier in the time series. Decoding for all comparisons except object decoding remained above chance until around 300ms, which included five subsequent stimulus presentations. There was no difference between distractor processing on boat target and star target trials (BF < 1/10) for all categorical contrasts.

Temporal generalisation analyses were performed to compare categorical decoding between the 5Hz and 20Hz conditions. For all three categorical levels, we observed similar onsets between presentation durations, but longer subsequent processing for the 5Hz condition relative to the 20Hz condition (Figure 6). Notably, for the animacy distinction there was no evidence of generalisation between the 5Hz sequence around 500-600ms and the 20Hz sequence at any time point, despite a difference between decoding accuracies during this time period (as was seen in Figure 5). This suggests that a high-level animacy-related process was present in the 5Hz condition but absent in the 20Hz condition. The temporal generalisation analyses also showed consistent below chance generalisation between the early and late responses. This phenomenon is consistent with previous decoding studies on visual object categorisation (Carlson et al., 2013; Cichy et al., 2014), and has been suggested to be caused by the stimulus offset, or by an adaptation or inhibition signal (Carlson et al., 2011, 2013; Contini et al., 2017).

**Figure 6.**
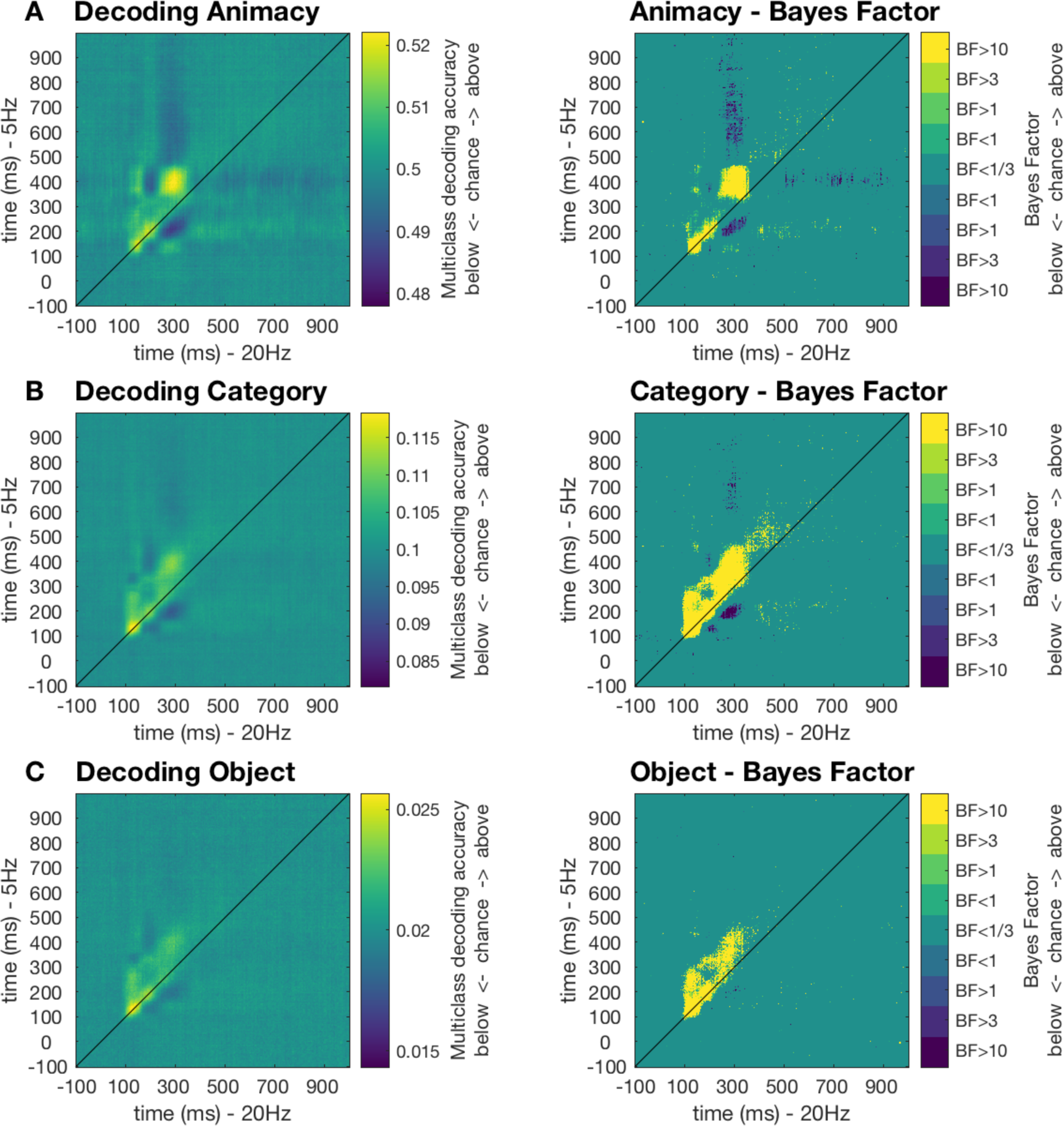
Temporal generalisation results. A) Decoding animacy (animate versus inanimate; chance = 50%). B) Decoding 10-way category (e.g., mammal, tools; chance = 10%). C) Decoding 50-way object categories (e.g., dog, giraffe; chance = 2%). The left columns show classifier generalisation performance for the three categorical levels between the different presentation durations. The right columns show corresponding thresholded Bayes Factors (Yellow indicating above chance, and blue indicating below chance decoding). Higher than chance generalisation (yellow) above the diagonal indicates slower processing in the 5Hz condition relative to the 20Hz condition.

### Representational dynamics of 200 stimuli

Emerging representational structures of the 200 stimuli were studied in the Representational Similarity Analysis (RSA) framework (Kriegeskorte & Kievit, 2013; Kriegeskorte, Mur, & Bandettini, 2008; Kriegeskorte, Mur, Ruff, et al., 2008). A neural representational dissimilarity matrix (neural RDM) was created for each subject, and each time point containing the dissimilarities between all 200 stimuli. Neural RDMs were modelled as a linear combination of six candidate models; low-level image silhouette, colour and luminance models, and one model for each of the three categorical levels. We then analysed the mean beta estimates of the candidate models (Figure 7). For both presentation rates, the silhouette model captured the early response in the data, followed by the colour, object, and category models. These results quantify the contribution of low-level visual features to neural dissimilarities. While low-level features were represented early in the signal, the categorical models also explained unique variance in the data. In the 5Hz condition, the animacy model emerged last, while in the 20Hz sequences the animacy model did not explain variance in the neural RDM at any time point. To visualise and qualitatively explore the dynamic representational structure, we created RDMs and a two-dimensional embedding of all 200 images from 5Hz and 20Hz sequences. Figure 8 shows these embeddings for 5Hz at two time points, 200 and 400ms, which are the time points where the category and animacy models were represented strongest in the signal (as observed in Figure 7). In these embeddings, the distance between images reflects their mean dissimilarity across subjects (Figure 8; see supplementary material for neural RDMs and two-dimensional embedding for 5Hz and 20Hz at all time points).

**Figure 7.**
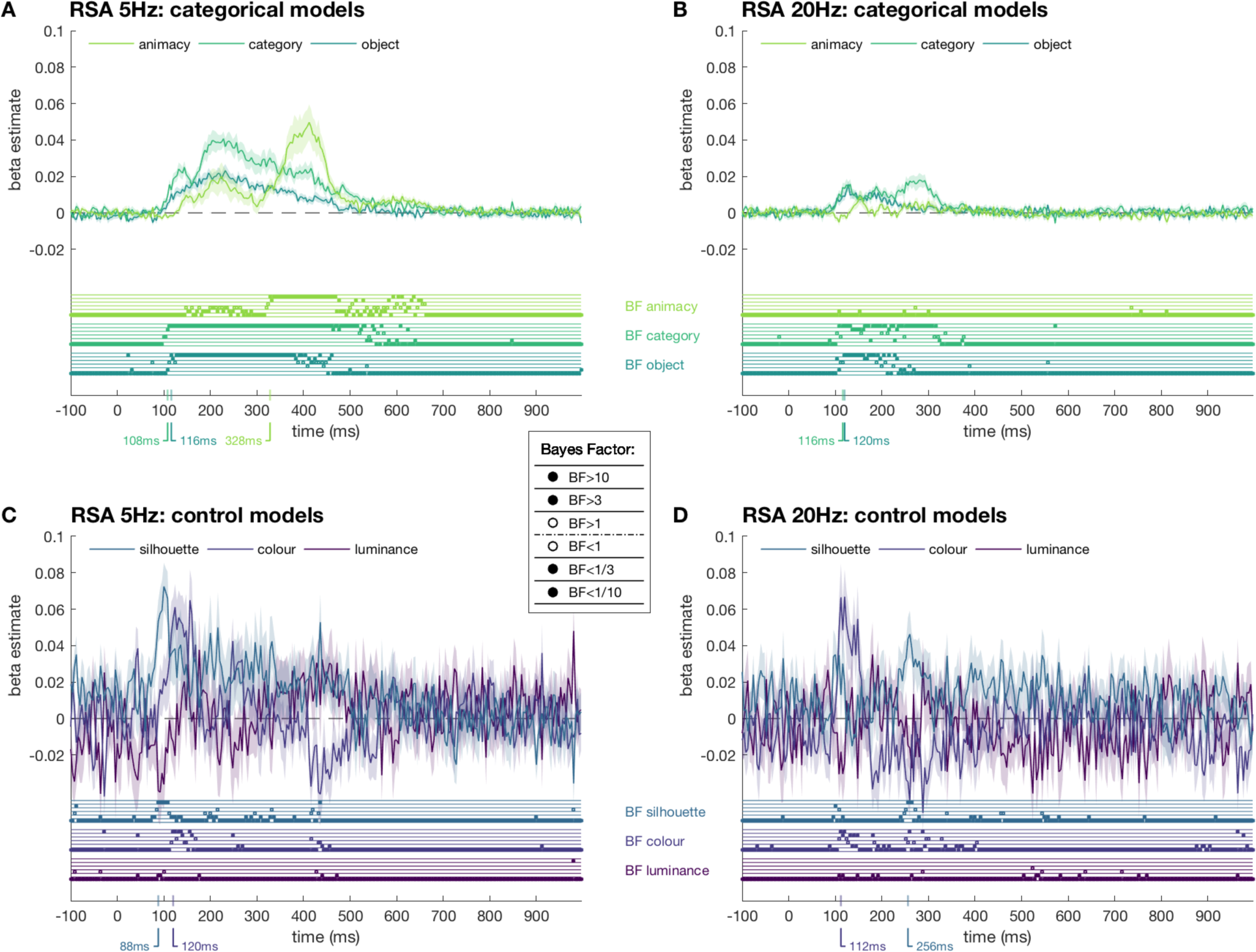
RSA model tests. A) 5Hz categorical models. B) 20Hz categorical models. C) 5Hz image feature control models. D) 20Hz image feature control models. The neural RDMs of each subject were modelled as linear combination of six candidate models: three categorical models and three image feature models. Lines show estimated betas for the models. Shaded areas reflect the standard error across subjects. Dots below plots indicate the thresholded Bayes Factors (BF, see inset) for each beta estimate. Annotated below the x-axis are the time points where the BF first exceeded 3 (for at least 2 consecutive time points).

**Figure 8.**
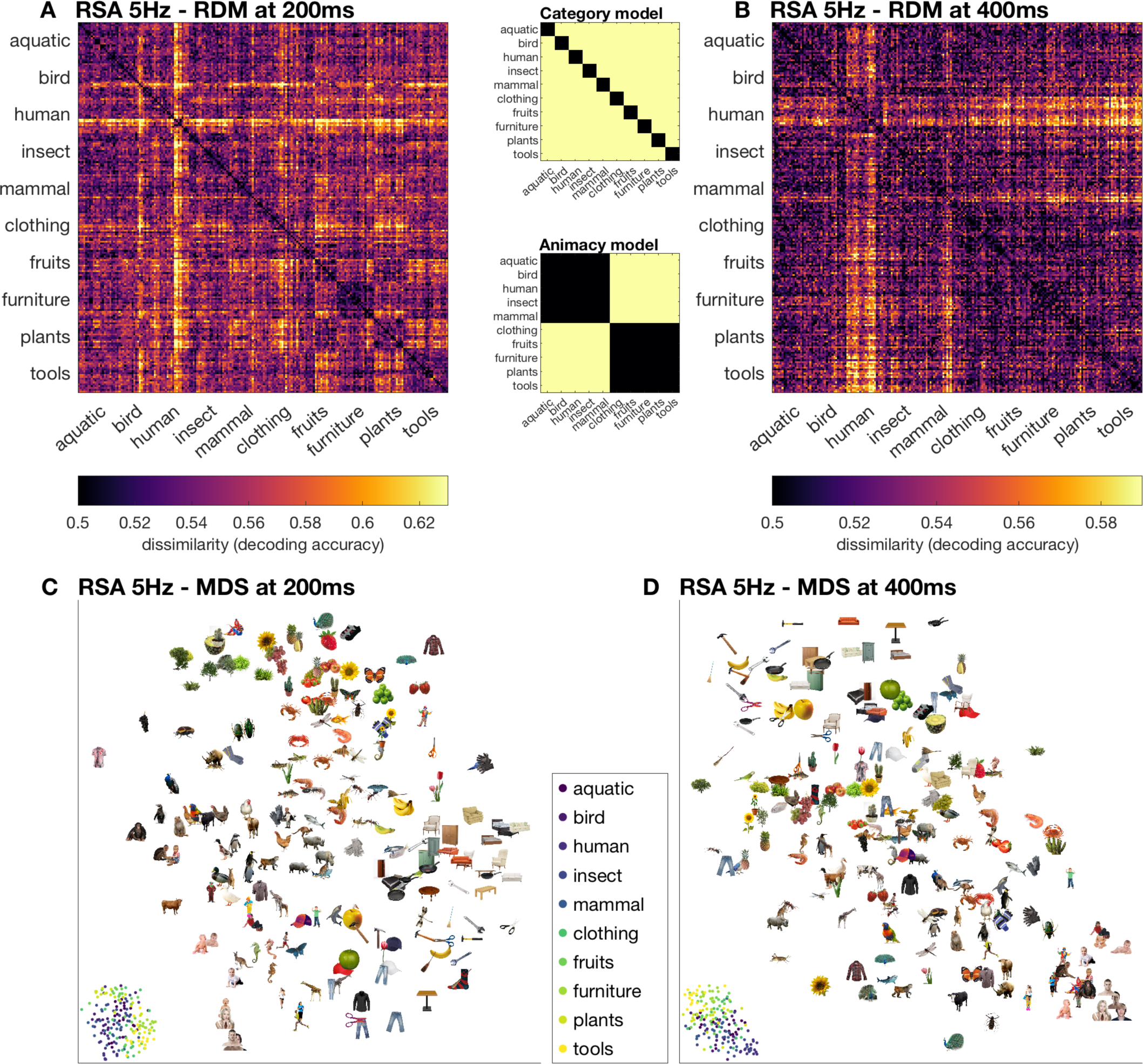
Representational structure of images in 5Hz sequences at two time points. A) Neural dissimilarity matrix at 200ms. B) Neural RDM at 400ms. The RSA model testing (Figure 7) showed that the structure at 200ms best resembles the category model and structure at 400ms best resembles the animacy model. C) Embedding of stimuli in a two-dimensional space reflects their pairwise distances at 200ms. D) Embedding at 400ms. Stimuli that are shown further apart in this representation evoked more dissimilar neural responses. In the bottom left corner of each plot, the same arrangement is shown, with images represented by dots coloured according to the 10 categories (see inset).

## Discussion

In the current study, we characterised the representational dynamics of a large number of images in fast presentation sequences. Previous work has used MEG and EEG decoding to investigate representations of much smaller image sets using slow image presentation paradigms (Carlson et al., 2013; Cichy et al., 2014; Contini et al., 2017; Grootswagers, Ritchie, et al., 2017; Kaiser et al., 2016; Kaneshiro et al., 2015; Proklova et al., 2017; Ritchie, Tovar, & Carlson, 2015; Simanova, van Gerven, Oostenveld, & Hagoort, 2010); here we extend this work by looking at the representations of 200 objects during RSVP using standard 64-channel EEG. For 5Hz and 20Hz sequences, all 200 images could be decoded at four different categorical levels. Furthermore, neural responses to targets were distinct from those to distractor stimuli. Above-chance decoding outlasted subsequent image presentations, supporting the idea that multiple object representations can co-exist in the visual system at different stages of processing (Marti & Dehaene, 2017). In keeping with the known hierarchical nature of the visual system, RSA model testing suggested neural responses relied on low-level visual features early in the time series, and subsequent processing was associated with increasing category abstraction (Carlson et al., 2013; Cichy et al., 2014). Overall, we show the unprecedented ability of the human brain to process images when pushing the limits of temporal perception.

Target decoding results revealed that neural responses to distractors diverged from star target responses much earlier than boat targets. This supports our initial hypothesis that star targets would be distinct from other images based on low-level visual features, unlike boat targets. The behavioural results, however, revealed target detection did not differ across boat and star trials, indicating that there was no “pop-out” effect of stars. This is despite anecdotal reports that participants found the star targets easier. Target versus distractor decoding for boats and stars peaked at 500ms, supporting previous evidence that high level cognitive processes mediate temporal selection (Marti & Dehaene, 2017; Sergent, Baillet, & Dehaene, 2005). These results suggest that distinguishable low-level features do not help with target detection in RSVP sequences, at least in the current design with such high variation in distractor images.

Target processing did not differ markedly across the different experimental durations. In both the 5Hz and 20Hz sequences, targets could be distinguished from distractors for a long period of time, but this was exaggerated for the 5Hz condition, where decoding was above chance for over 1000ms, compared to 800ms in the 20Hz condition. Decoding was also higher in the 5Hz condition relative to the 20Hz condition, but the dynamics of temporal selection processes were largely the same. The time of peak decoding (500ms) was the same for both conditions, and time generalisation analyses revealed neural processes occurred at the same latency in both conditions. This suggests that processes of target selection are largely the same regardless of image presentation duration and frequency. Notably, target processing was much more prolonged than categorical decoding for distractors, again indicative of higher level cognitive processes at play for target detection. Note that the current experimental design did not allow us to see which targets in the stream were missed, but effects are likely to be amplified for correctly detected targets. Indeed, Marti & Dehaene (2017) found that late responses were sustained for reported stimuli. Taken together, our results show that late target-related responses do not differ dramatically in faster sequences relative to slower sequences.

Neural responses to the 200 non-task-relevant (distractor) objects are indicative of fairly automatic early visual processing and divergence at later processing stages according to image duration. For all contrasts, image presentation duration and cognitive task set did not influence the earliest processing stages. When looking at decoding for the durations separately, onsets seemed to be earlier for the 5Hz than 20Hz conditions, in accordance with recent work showing earlier onsets for longer image durations (Mohsenzadeh et al., 2018). It is important to note, however, that higher signal strengths can also lead to earlier decoding onsets (Grootswagers, Wardle, et al., 2017), thus differences between onsets must be interpreted with caution in the context of larger peak decoding. Crucially, here Bayes factors revealed evidence for no difference in decoding at these early time points between the 5Hz and 20Hz image sequences (<150ms from image onset). Results from the temporal generalisation approach supported this view, by showing that initial processing stages occurred at the same time for the 5Hz and 20Hz sequences, as seen by the above-chance decoding on the diagonal in Figure 5. Finally, for the three categorical levels (animacy, category and object), Bayesian analyses revealed distractor processing did not differ between boat and star trials. These results suggest that initial neural responses to all visual stimuli were similar regardless of their presentation duration.

Previous work has shown that, using MEG, it is possible to use decoding to investigate target-related processes in RSVP streams (Marti & Dehaene, 2017; Mohsenzadeh et al., 2018a). For example, Mohsenzadeh et al. used 306-channel MEG to decode 12 target faces from 12 non-target objects in RSVP streams, analysing only the middle image in the stream to study feedforward versus feedback processes. As part of a study investigating temporal selection mechanisms, Marti & Dehaene showed that a classifier trained on 5 categories using a separate localiser could generalise to distractor items around the target. In contrast to these studies, here we decoded object representations using a 64-channel EEG, a much larger set of images (200) in a sequence, and no separate localiser. The results from our approach also corroborated previous work decoding the representations of objects presented in isolation (Carlson et al., 2013; Cichy et al., 2014; Kaneshiro et al., 2015). Our results showed that decoding objects in RSVP streams have similar decoding onsets as previously reported (Carlson et al., 2013; Cichy et al., 2014; Kaneshiro et al., 2015). This validates the RSVP approach as a method to study representational dynamics. We further found that the 20Hz condition limited visual processing compared to 5Hz, which shows that this paradigm can be utilised to bias the extent of visual processing at different image presentation rates. In sum, our results confirm that long ISIs are not necessary for multivariate analyses. This thus allows analysing all presentations in an RSVP sequence, rather than limiting the scope to selected presentations (e.g., targets) in the streams. Here we have demonstrated the potential by studying the representational dynamics of 200 objects in one short EEG session. Future work can adopt similar approaches to investigate for example prediction, priming, masking, or attentional effects on the processing of distractors in RSVP sequences.

Despite similar early processing stages, later processing diverged according to image presentation duration. Representations during 5Hz sequences were stronger and lasted longer than those during 20Hz sequences, and temporal generalisation analyses showed that processes were prolonged for the 5Hz relative to the 20Hz condition. It could be that longer image durations allow more consolidation, potentially due to recurrent processing. It is also possible that longer durations allow time to reach some kind of threshold, which triggers further processing. Note that image duration and ISI are conflated in this design, so we cannot conclude whether or if stronger and longer processing occurs due to longer image presentation or due to delayed masking from the next stimulus. Future work can build on this approach to investigate the temporal limits of visual perception.

The RSA regression analyses provided insight into the differences in processing between the 5Hz and 20Hz sequences. The category decoding analyses were performed using a leave-one-exemplar out cross-validation approach, which means that the classifier always had to generalise to new images, reducing the likelihood that low-level features would drive the results. However, there can still be consistent low-level features between the categories that can contribute to classification. The regression RSA technique aimed to dissociate the unique contributions of each of the categorical and low-level featural models. In accordance with the decoding results, processes early in the time series (∼100-150ms) were mostly explained by the low-level silhouette model and then the colour model for the 5Hz and 20Hz conditions (Carlson et al., 2011). Subsequent processing, however, elucidated the differential contributions of the different categorical contrasts, and how this varied for the different image durations. For the 5Hz condition, the category model appeared to have the largest unique contribution around 200ms, and the animacy model accounted for the most variance at about 400ms, indicating that increasing category abstraction occurred at higher levels of visual processing (Carlson et al., 2013; Cichy et al., 2014; Contini et al., 2017; Kriegeskorte, Mur, Ruff, et al., 2008). In contrast, the animacy model had no unique contribution to the signal for the in 20Hz sequences. The time course of the animacy model regression for the 5Hz condition (>350ms) suggests that the animate-inanimate difference might exclusively account for the prolonged decoding in the 5Hz condition relative to the 20Hz condition. This could imply that a high-level animacy effect requires sufficient evidence accumulation to proceed, which does not happen at 20Hz presentation rates. The finding that longer image presentations allow higher level processing is supported by steady-state visual evoked potential (SSVEP) work showing that images presented at faster frequencies are biased towards earlier visual processes in contrast to slower frequencies which allow higher level processing (Collins, Robinson, & Behrmann, 2018).

When qualitatively inspecting the visualisation of the representational structure (Figure 8), we noticed a clear categorical organisation in the 5Hz presentation condition. At 200ms in the response, the structure reflected mostly natural versus artificial, with plants, fruits and animals all clustering on one side (Figure 8C). In line with the decoding and RSA results, the structure at 400ms showed a clear animate – inanimate distinction (Figure 8D) (Caramazza & Shelton, 1998), which is commonly observed in neural responses in the ventral temporal cortex (Cichy et al., 2014; Konkle & Caramazza, 2013; Kriegeskorte, Mur, Ruff, et al., 2008; Proklova, Kaiser, & Peelen, 2016) and has been shown to match human categorisation behaviour well (Bracci & Op de Beeck, 2016; Carlson, Ritchie, Kriegeskorte, Durvasula, & Ma, 2014; Grootswagers, Cichy, & Carlson, 2018; Mur et al., 2013; Ritchie, Tovar, & Carlson, 2015). In the animate – inanimate organisation primates were located at the far end of the animate side, which may reflect a continuum of biological classes in the brain (Connolly et al., 2012; Sha et al., 2015) or typicality (Grootswagers, Ritchie, et al., 2017; Iordan, Greene, Beck, & Fei-Fei, 2016; Posner & Keele, 1968; E. H. Rosch, 1973; E. Rosch & Mervis, 1975). No animacy structure was apparent for the 20Hz condition (as evidenced by the RSA results), but rather individual categorical clusters seem to have emerged (in line with the RSA results), such as human faces, and, later, humans and primates as a category (see Supplementary Material). Interestingly, in these visualisations gloves were grouped with humans and primates, which could mean they were perceived as body parts, rather than inanimate objects. While these visualisations allow for such qualitative speculation, the quantitive RSA modelling results highlight the level of detail in the representation structure that can be obtained using EEG decoding and fast presentation rates. Here, we used a common 64-channel EEG, but future work can use this approach in combination with high-density EEG or other neuroimaging methods that are sensitive to finer spatial patterns, such as MEG.

One remaining question is the role that low-level image statistics play in our results. The RSA approach showed that low-level control models explained early neural responses to the stimuli. The current stimulus set consisted of segmented coloured objects, which were not matched on low-level features such as colour, orientation, shape, and size. Future work can build on the current paradigm using a stimulus set that for example contains orthogonal shape and category dimensions (Bracci, Kalfas, & Op de Beeck, 2017; Bracci & Op de Beeck, 2016; Proklova et al., 2017, 2016), or test the decodability of these features using for example texture stimuli with similar features (Long, Konkle, Cohen, & Alvarez, 2016; Long, Yu, & Konkle, 2017). Such extensions can help unravel the relationship between object features and categories, and increase our understanding of how this inherent relationship guides categorical abstraction in the visual system.

In conclusion, our results show that we can study the representational dynamics of more than 200 objects in one short EEG session. We were able to characterise the time courses of multiple categorical contrasts from the same images, indicating that all objects reached abstract categorical stages of perception despite being presented for short durations. Here, we took advantage of the high temporal resolution of both the human visual system and common neuroimaging techniques such as EEG and MEG. These results confirm that long ISIs are not necessary for multivariate analyses, as they do not require a resting baseline as in ERP analyses. Thus, future MVPA studies on visual perception should consider using fast presentation rates as this allows for a substantial increase of the number of presentations, stimuli, or experimental conditions. This offers unprecedented potential for studying the temporal dynamics of visual perception and attention.

## Acknowledgements

We thank Nick McNair for assisting with data collection. This research was supported by an Australian Research Council Future Fellowship (FT120100816) and an Australian Research Council Discovery project (DP160101300) awarded to T.A.C. The authors acknowledge the University of Sydney HPC service for providing High Performance Computing resources. The authors declare no competing financial interests.

