## Supplementary material for "The representational dynamics of visual objects in rapid serial visual processing streams" (from another version of this preprint: version 3, 2018-11-26)

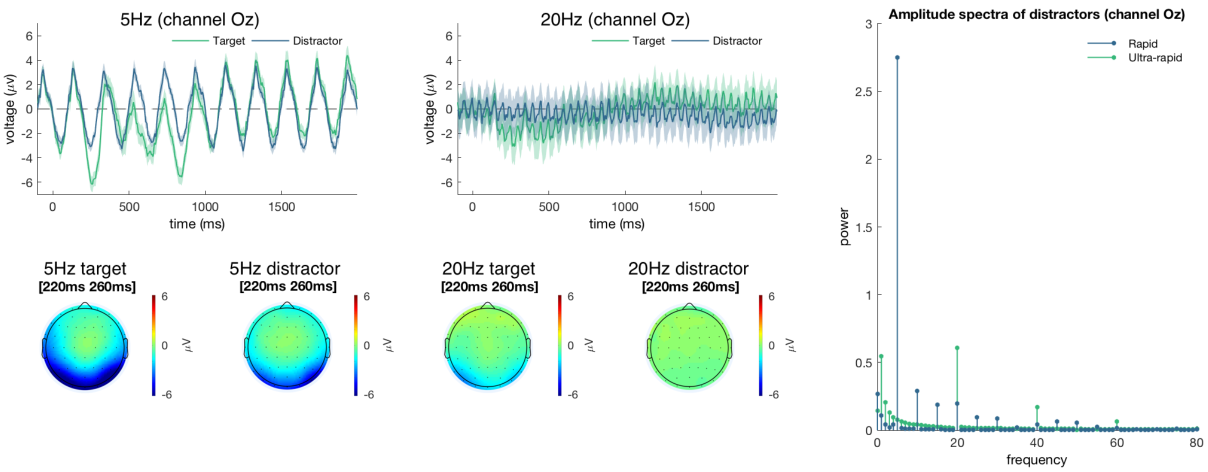


**Figure S1.** Event-related potentials for targets and distractors, frequency response for distractors, and topoplots for the 5Hz and 20Hz sequences.

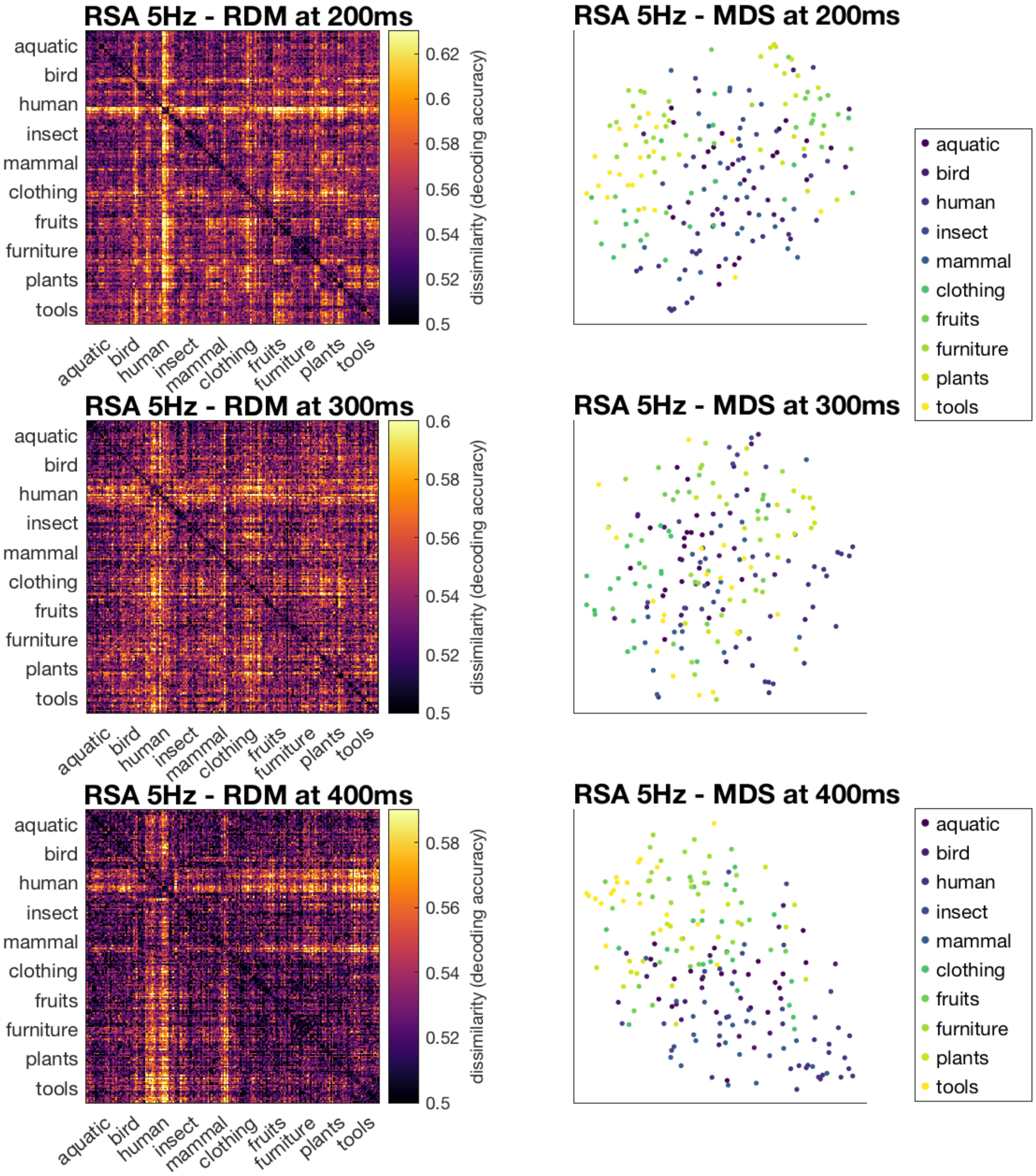


**Figure S2.** 5Hz Neural RDMs and MDS solutions for three time points corresponding to the visualisations in Figure 8.

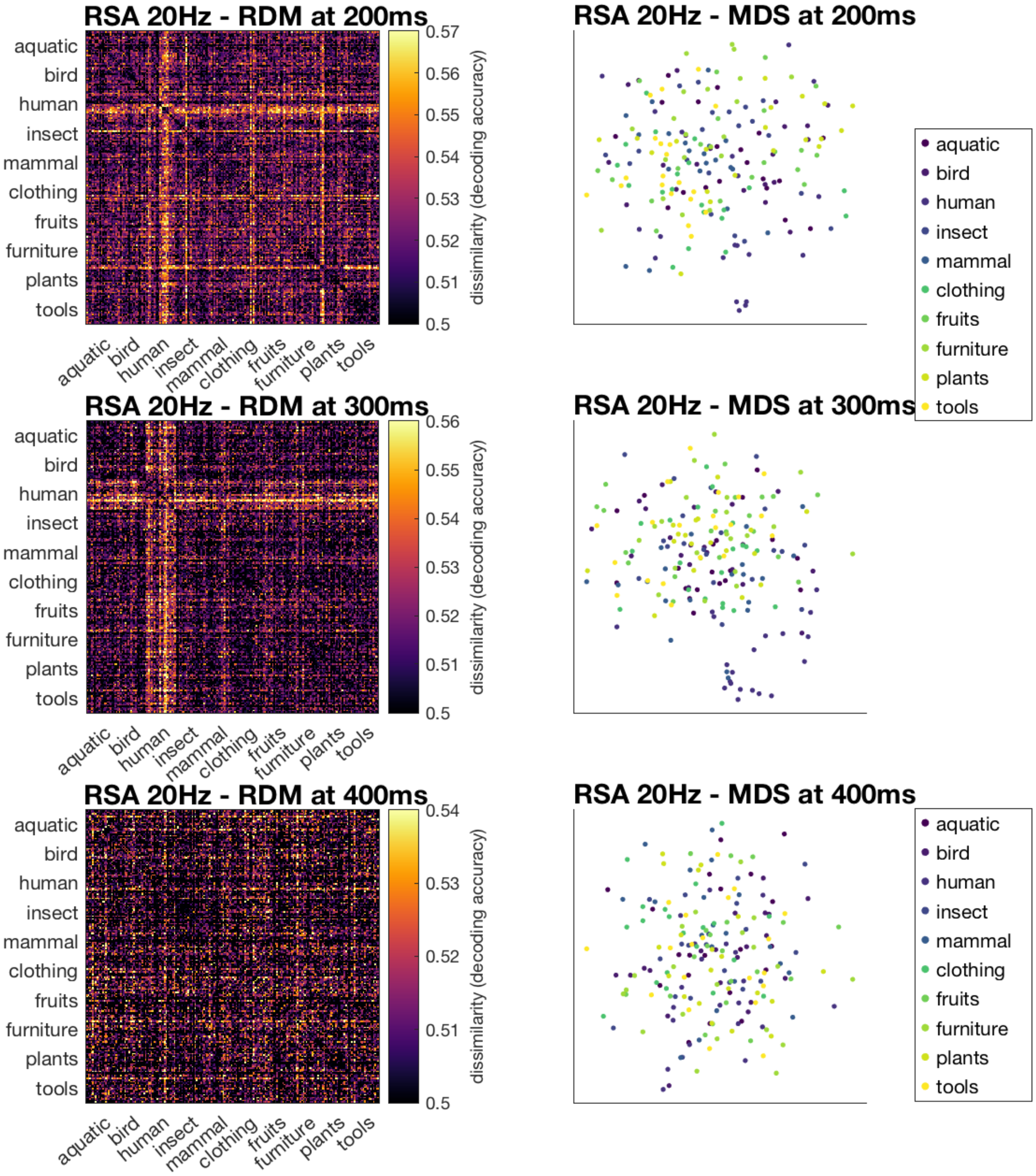


**Figure S3.** 20Hz Neural RDMs and MDS solutions for three time points corresponding to the visualisations in Figure 8.
